# Germline transformation of the West Nile Virus and avian malaria vector *Culex quinquefasciatus* Say using the piggyBac transposon system

**DOI:** 10.1101/2023.12.01.569580

**Authors:** Katherine Nevard, Rajdeep Kaur, Tim Harvey-Samuel

**Affiliations:** The Pirbright Institute, Ash Road, Pirbright, Surrey, GU24 0NF

**Keywords:** *Culex quinquefasciatus*, piggyBac, genetic engineering, mosquito control, genetic control

## Abstract

*Culex quinquefasciatus* Say is a mosquito which acts as a vector for numerous diseases including West Nile Virus, lymphatic filariasis and avian malaria, over a broad geographical range. As the effectiveness of insecticidal mosquito control methods declines, the need has grown to develop genetic control methods to curb the spread of disease. The piggyBac transposon system - the most widely used genetic transformation tool in insects, including mosquitoes - generates quasi-random insertions of donor DNA into the host genome. However, despite the broad reported species range of piggyBac, previous attempts to use this tool to transform *Culex quinquefasciatus* mosquitoes have failed. Here we report the first successful transformation of *Culex quinquefasciatus* with the piggyBac transposon system. Using commercially synthesised piggyBac mRNA as a transposase source, we were able to generate three independent insertions of a *ZsGreen* fluorescent marker gene, with transformation efficiencies of up to 5%. Through this work, we have expanded the genetic toolkit available for the genetic manipulation of *Culex* mosquitoes and thus removed a barrier to developing novel genetic control methods in this important disease vector.

## 1. Introduction

*Culex quinquefasciatus* Say is a mosquito in the *Culex pipiens* species complex, which acts as a vector for a wide range of human and wildlife pathogens across tropical and subtropical regions (LaPointe, 2012). As a vector of avian malaria, *Cx. quinquefasciatus* has devastated populations of endemic bird species on islands including Hawaii and New Zealand (Harvey-Samuel et al., 2021), with Hawaiian honeycreepers being particularly susceptible to the disease (Atkinson et al., 2000, 1995). Furthermore, in Hawaii the threat to avifauna has grown as climate change has driven an increase in prevalence of malaria infection in higher elevation areas that previously served as refuges from the disease (Atkinson et al., 2014; Neddermeyer et al., 2023). Birds are the primary hosts for *Cx. quinquefasciatus*, however they feed on a wide variety of hosts including humans, spreading diseases such as West Nile Virus, lymphatic filariasis and Western equine encephalitis (LaPointe, 2012). Lymphatic filariasis, caused by infection with the nematode worm, *Wucheria bancrofti*, causes severe swelling leading to pain and disability. Efforts to control the disease have incurred great costs, with over 9 billion treatments administered since the year 2000 (WHO, 2023). West Nile Virus is the leading cause of domestically acquired arboviral disease in the USA (Soto et al., 2022, CDC, 2023) and has been attributed to $56 million per year in healthcare costs (Shankar et al., 2014). As the effectiveness of insecticidal control of this mosquito declines due to increasing insecticide resistance (Delannay et al., 2018; Pocquet et al., 2013; Talipouo et al., 2021) interest has grown in developing transgene-based ‘genetic control methods’, which additionally offer the ecological benefit of being targeted towards a single species (Wang et al., 2021). Proposed designs have included ‘First generation’ genetic control systems such as fsRIDL which function analogously to the classical Sterile Insect Technique, relying on mass-releases of insects carrying dominant, repressible, lethal genes to suppress the reproductive potential of wild populations. And ‘second generation’ technologies also known as gene drives, aiming to drive these lethality (or other) traits through a population, e.g. by engineering super-mendelian inheritance, thereby negating the need for the expensive mass-release systems of first generation technologies (Siddall et al., 2022).

Regardless of the system in question, all these genetic control technologies require a method for engineering the genome of their target species. While gene-drive systems often make use of the site-specific CRISPR/Cas9 genome editing technologies – where exogenous DNA sequences are introduced into the pest genome at a precise point through the homology-directed repair (HDR) pathway - first generation technologies, and insect genetic engineering more broadly, has relied on transposon-based methods which create quasi-random integration events. An advantage of these quasi-random integrations is that they provide a ‘brute-force’ method for overcoming the influence of surrounding genomic sequences on the function (i.e. expression levels) of an integrated transgene (positional effects). Here, multiple independent integrations can be made with no *a-priori* knowledge of how the genomic context of those integrations will influence the transgene. The function of the transgene can subsequently be tested in those multiple independent insertions, and the insertion with the most suitable expression levels (and therefore genomic context) for a particular application taken forward.

The most widely used transposon-system for insect genetic engineering is piggyBac (PB). PB is a Class II (cut and paste) transposable element that was originally discovered in the cabbage looper moth, *Trichoplusia ni*, (Fraser et al., 1983) and was subsequently adapted for use as a transformation tool (Handler et al., 1998). Transposition occurs at TTAA sites (Wang and Fraser, 1993) and is therefore relatively unconstrained, generating effectively random insertions in the genome, which together with its high efficiency (Eckermann et al., 2018), capability of precise excision (Elick et al., 1996), and large cargo size (Li et al., 2011), has led to the widespread utilisation of PB for insect transgenesis (Gregory et al., 2016). Transgenic *Cx. quinquefasciatus* have previously been generated using CRISPR-Cas9 mediated HDR (Feng et al., 2021; Harvey-Samuel et al., 2023; Purusothaman et al., 2021) and the Hermes transposable element (Allen et al., 2001; Allen and Christensen, 2004), however to date there are no reports of successful transformation using the PB system. This is surprising, given the broad species range, including yeast, plants, insects and mammals (Ding et al., 2005; Gregory et al., 2016; Nishizawa-Yokoi and Toki, 2021; Zhu et al., 2018), and previous success using PB in other mosquito genera (Grossman et al., 2001; Labbé et al., 2010a; Lobo et al., 2002; Nolan et al., 2002; Perera et al., 2002). Indeed, published, unsuccessful, attempts at generating PB insertions in *Cx. quinquefasciatus*, using both plasmid and *in vitro* transcribed mRNA transposase sources suggested that this insect may prove more difficult than other species to transform using this system (Anderson et al., 2019).

Here, we attempt to overcome this limitation by translating findings we initially observed in the diamondback moth (*Plutella xylostella*) to *Cx. quinquefasciatus*. The diamondback moth is a model for lepidopteran genetic control tools and is a species in which PB has been widely used. Despite this, transformation rates in this species are relatively low (Gregory et al., 2016). Previously, we observed in diamondback moth that transformation efficiencies could be improved by a factor of 10 or more by using commercially synthesised PB mRNA as a transposase source (1.0%, 2.4%), in comparison to codon optimised in-vitro transcribed PB mRNA (0.1%, 0.7%) (Xu et al., 2022) or plasmid-encoded transposase (0.48%-0.68%) (Martins et al., 2012). To explore whether this approach would be viable in *Culex* mosquitoes, we set out to generate PB transformants in *Cx. quinquefasciatus* using the same commercially synthesised PB transposase mRNA. With this, we hoped to expand the genetic manipulation toolkit available in the *Culex* genus and remove a barrier to progress in the field of genetic control for this widespread and important disease vector.

## 2. Methods

### 2.1 Mosquito rearing

*Culex quinquefasciatus* adults were maintained in cages (Bugdorm, Taiwan), kept at 27°C, 70% humidity and on 12/12 hour light/dark cycle with one hour of dawn and dusk. Adult mosquitoes were provided with 10% sucrose solution *ad libitum*. Females were fed with defibrinated horse blood (TCS Biosciences, Buckingham, UK) through a membrane feeder (Hemotek, Blackburn, UK) placed on top of the cage with the reservoir covered with a piece of hog gut (sausage casing). Egg rafts were collected in 2oz portion cups, half filled with water, that were placed inside the cage two days after bloodfeeding. Larvae were fed with fish food pellets (Su-Bridge Pet Supplies Ltd, Thetford, UK).

### 2.2 Injection mix

The *piggyBac* transposase open reading frame (Supplementary Text 1) was synthesised as purified mRNA, supplied by tebubio (Le Perray-en-Yvelines, France). In brief, this was capped using CleanCap AG Polyadenylated (120A), treated with DNase and phosphatase, silica membrane purified and received as a solution in 1mM sodium citrate, pH6.4. This was diluted upon receipt to 880ng/l with nuclease free water and stored at -80°C. The injection mix was made up of *piggyBac* mRNA (440ng/ul) and plasmid PL1127 (300ng/ul) which contained a *ZsGreen* marker gene under control of the Hr5/IE1 promotor, flanked by *piggyBac* inverted terminal repeats. (Genbank accession number: OR883676; Fig 1). The mix was centrifuged before use at 11,000 rpm for 10 minutes, then kept on ice throughout the injections.

**Figure 1.**
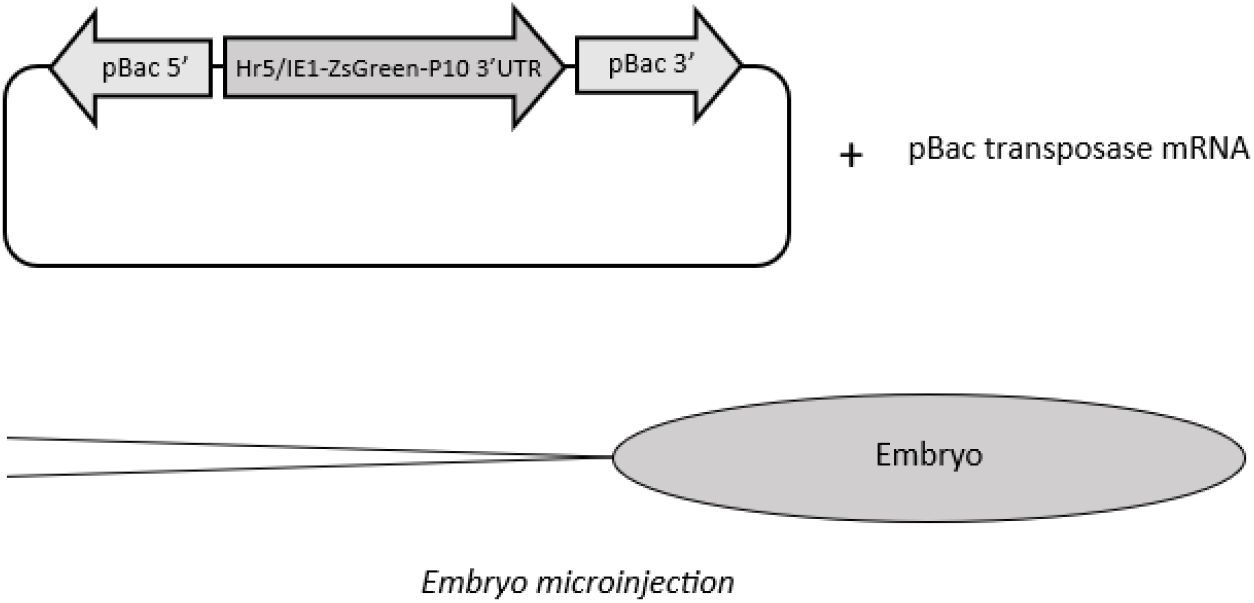
Depiction of microinjection mix components and microinjection process. PL1127 plasmid is shown on the left, containing Hr5/IE1 promotor sequence, *ZsGreen* marker and P10 3’ UTR, with piggyBac flanks (not drawn to scale). PL1127 was injected with piggyBac transposase mRNA into mosquito embryos.

### 2.3 Embryo microinjections

Embryo microinjections were performed as described by (Anderson et al., 2019) with the following modifications. Needles were pulled on Sutter P2000 laser based micro-pipette needle puller (Sutter Instruments, Novato, CA, USA), using the following settings: Heat = 715, FIL = 4, VEL = 40, DEL = 128, PUL = 134. Once injected, the injection slides were rinsed to remove oil then submerged in water, egg side up, in a petri dish and left for approximately 24 hours. The slides were then transferred to a dH_2_0 filled tray, placed egg side down on the surface of the water, where hatching could occur. Once hatched, the generation 0 (G0) L1 larvae were transferred to another tray. This transferral process was intended to minimise any deleterious effects of halocarbon oil exposure.

### 2.4 Crosses and screening

G0 pupae were sexed and screened for transient expression of the *ZsGreen* marker under a Leica MZ165FC microscope (Leica Biosystems, Milton-Keynes UK). Individuals showing transient expression were recorded (Supplementary Table 1) but treated the same as non-transient expression individuals. Groups of up to 10 males or 10 females were put into separate cages. Males were crossed with WT females in a 1:5 ratio. Females were crossed with WT males in an approximately 1:1 ratio. Each separate cage of crossed males or females was deemed a separate pool; each given a letter to denote the pool name. Cages were blood fed three days after crossing and egg cups were added two days after the blood feed. Progeny from these cages were termed G1 and subsequent generations were denoted in the same format with ascending numbers. G1 eggs were collected, hatched and reared to L4 stage. L4 larvae were screened for *ZsGreen* fluorescence. This was repeated through a maximum of three ovipositions. Larvae identified as positive for the *ZsGreen* marker were reared to adults and crossed individually with WT in the same ratios as described previously. Photographs of transgenic individuals were taken with a Leica DFC camera.

### 2.5 Location of insertions

G1 adults and some additional G3 adults were frozen individually after mating/ovipositing. Genomic DNA was extracted using Nucleospin Tissue Kit (Macheray-Nigel, Düren, Germany). Adaptor ligation-mediated PCR was performed according to a published method (O’Malley et al., 2007) which aims to allow location of PB insertions through sequencing of the genomic regions flanking the PB insert. The exact procedure followed previous efforts in other insects (Martins et al., 2012).

## 3. Results

Two sets of microinjections were performed on *Cx. quinquefasciatus* embryos. In the first set, 145 eggs were injected and 60 survived to adult stage (41.3%) (Table 1). In the second set, 99 embryos were injected and 40 survived to adults (40.4%). Of the 100 survivors, 50 females were grouped into five pools and 50 males were grouped into five pools that were crossed to wild type mosquitoes in the aforementioned ratios. Larvae were screened from the following generation (G1) for the presence of the marker gene, through a maximum of three ovipositions. Larvae expressing the marker gene were identified in three of the 10 pools. The minimum integration rate from both sets of injections combined was 3.0%, calculated as follows:

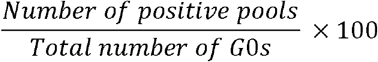

This calculation represents a minimum integration rate, since G0 mosquitoes were pooled into groups of 10, and thus only one transposition event would be counted per pool, even if more were to occur. When calculated individually for the first and second set of injections, the minimum integration rates were 1.7% and 5.0% respectively.

**Table 1.** Survival and integration rates for the two sets of microinjections performed to generate PL1127 transgenic lines.

| **Eggs**<br>**injected** | **Adult G0**<br>**survivors** | **Adult G0**<br>**survival (%)** | **G1s**<br>**screened** | **Transgenic**<br>**pools**<br>**identified** | **Minimum**<br>**integration**<br>**rate (%)** |
| --- | --- | --- | --- | --- | --- |
| 145 | 60 | 41.3 | 7102 | 1 | 1.7 |
| 99 | 40 | 40.4 | 2637 | 2 | 5.0 |

Transgenic mosquitoes were checked for marker visibility at L4 larval, pupal and adult stages and were shown to exhibit bright fluorescence at all of these stages (Fig 2). Of the G0 pupae that were screened for transient expression of the marker gene, 44.6% and 71.0% showed expression for the first and second sets on injections respectively. A small subset of the slowest developing individuals from each set of injections were not screened for transient expression, as they were separated as larvae and allowed to pupate and eclose over the weekend.

**Figure 2.**
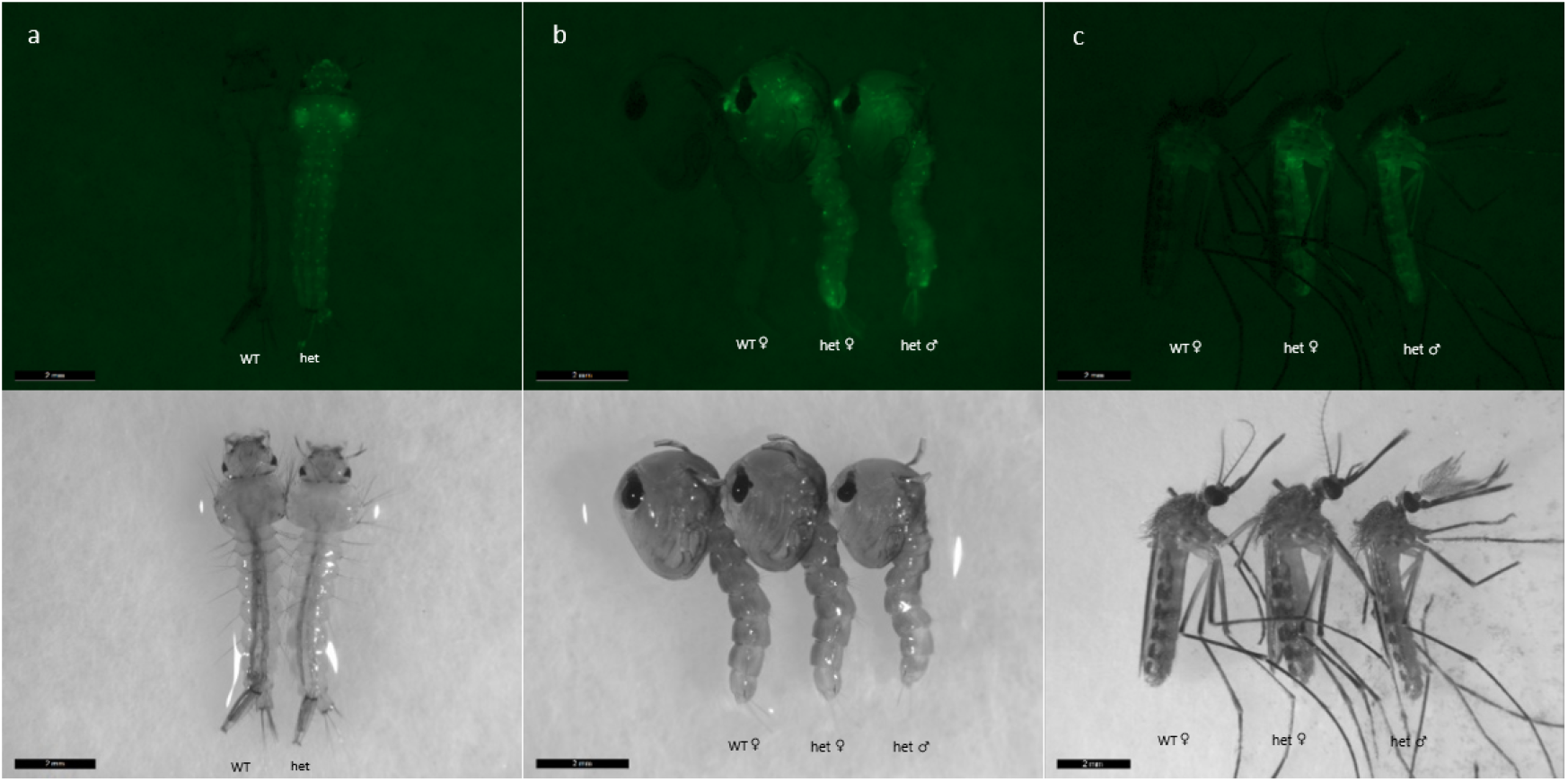
Photomicrographs of PL1127 mosquitoes from line K1 at a) larval, b) pupal and c) adult life stages, next to wild type (WT) for comparison. Images on top row are taken under fluorescent light to show *ZsGreen* fluorescent phenotype, and bottom row taken under white light. Images taken on Leica MZ165FC microscope with DFC365FX camera.

A total of five positive G1s survived to adult stage and were crossed individually with WT to generate G2 progeny; one from each of pools A and H, and three from pool K, which were renamed as pools K1, K2 and K3. G2s were screened at larval stage to assess the inheritance rate of the marker gene. A higher than 50% inheritance rate of the marker may have indicated multiple insertions of the gene. Pool H, K2 and K3 failed to produce any G2 progeny after multiple rounds of blood feeding. Exactly 50.0% of progeny from pool A were positive for the marker, and 54.8% of progeny from pool K1 were positive for the marker (Supplementary Table 2), supporting that these were likely single insertions.

5’ and 3’ genomic regions flanking the transgene insertion were identified for lines H and A. Blastn analysis of these concatenated sequences (5’ + 3’, separated by the TTAA insertion site) in the *Cx. quinquefasciatus* JHB reference genome showed unbroken, highly significant, matches across the query, providing further support that these sequences represented the insertion sites. Interestingly, each of the insertion sites was highly repeated across the genome (>100x), occurring on all three chromosomes, suggesting that each of these insertion sites occurred in multi-locus repeats. For Line K, despite exposing the gDNA sample to multiple restriction enzymes with differing recognition sites, only the 5’ flanking sequence could be identified. This flanking sequence could not be significantly matched to a genomic sequence, likely due to its short length. Sequences and details of genomic matches are provided in the Supplementary Materials (Supplementary Text 2).

## 4. Discussion

Prior to this study, transformation in *Culex* mosquitoes had only been achieved using the Hermes transposable element and CRISPR-based HDR. Hermes transformation was attained using plasmid transposase and saw a high transformation efficiency of 11.8%, generating two transgenic lines from 17 fertile individuals (Allen et al., 2001). A separate attempt yielded two egg rafts containing transgenics from a total of 32 female G0s (Allen & Christensen, 2004), which would give a rate of 6.25%. Our rates of 1.7% and 5% (mean 3%) were calculated from both male and female pools of individuals, including those that may not have been fertile, and thus represented a minimum rate. Through HDR using a plasmid donor, (Purusothaman et al. 2021) achieved a minimum integration rate of 1.6%, or 2.8% discounting the initial unsuccessful round of injections. 985 eggs were injected to produce a total of two transgenic lines. Feng et al. (2021), also produced transgenic *Cx. quinquefaciatu*s lines by HDR, achieving efficiency rates of 0%, 1.82% and 4.17% from the three constructs they injected. Through using PB as opposed to HDR, we were able to generate three transgenic lines from only 244 injected eggs, with the highest efficiency from a single set of injections being 5%. The insertions generated were in three independent positions within the genome. To generate these integrations at different genomic sites using HDR would require the design and testing of multiple guide RNAs and plasmid flanking sequences, as well as multiple sets of injections and would therefore be considerably less efficient.

PB is a widely used transformation tool in insects, with most recorded transformation rates within the range of 1-10% (Gregory et al., 2016). High transformation rates have been observed in *Aedes fluviatilis* (27%) (Rodrigues et al., 2006), *Anopheles albimanus* (43%) (Perera et al., 2002), and *Anopheles sinensis* (21.43%) (Liu et al., 2021), calculated based on the number of transgenic lines generated from fertile adults only. Average transformation efficiency in *Ae. aegypti* has been reported as lower, at 8% (Nimmo et al., 2006), with a later study finding a mean of 5.9% from a large Oxitec dataset (Gregory et al., 2016). The first PB integration of *Aedes albopictus* reported a minimum rate of 1%, adjusted to 2.2-3.6% when a 30-50% fertility rate was assumed (Labbé et al., 2010). Though our rate represents a minimum, with fertility not accounted for, our average rate of 3% still falls within the range previously observed in other mosquito species. If we were to also assume a 30-50% fertility rate, this would give a transformation efficiency of 4.3-6%.

Here we have demonstrated that PB transformation is possible in *Culex* mosquitoes and can achieve higher transformation rates than HDR. The benefit of generating effectively random insertions will allow for efficient testing and identification of different insertion sites to facilitate the optimisation of expression levels of inserted genes; an important aspect of generating genetic control tools. We anticipate these new findings will help to drive forward the development of novel genetic control methods within the *Culex* genus.

## Author Contributions

Experiments were performed by KN and RK. All authors contributed to experimental design and manuscript preparation.

## Data availability

All data is available within the manuscript.

## Acknowledgements

This work was supported by UK Biotechnology and Biological Sciences Research Council [BBS/E/I/00007033, BBS/E/I/00007038, and BBS/E/I/00007039] strategic funding to The Pirbright Institute.

