## Supplementary Material for "Germline transformation of the West Nile Virus and avian malaria vector *Culex quinquefasciatus* Say using the piggyBac transposon system"

**Supplementary Text 1.**

**Piggybac open reading frame**

ATGGGTAGTTCTTTAGACGATGAGCATATCCTCTCTGCTCTTCTGCAAAGCGATGACGAGCTTGTTGGTGAGGATTCTGACAGTGAAATATCAGATCACGTAAGTGAAGATGACGTCCAGAGCGATACAGAAGAAGCGTTTATAGATGAGGTACATGAAGTGCAGCCAACGTCAAGCGGTAGTGAAATATTAGACGAACAAAATGTTATTGAACAACCAGGTTCTTCATTGGCTTCTAACAGAATCTTGACCTTGCCACAGAGGACTATTAGAGGTAAGAATAAACATTGTTGGTCAACTTCAAAGTCCACGAGGCGTAGCCGAGTCTCTGCACTGAACATTGTCAGATCTCAAAGAGGTCCGACGCGTATGTGCCGCAATATATATGACCCACTTTTATGCTTCAAACTATTTTTTACTGATGAGATAATTTCGGAAATTGTAAAATGGACAAATGCTGAGATATCATTGAAACGTCGGGAATCTATGACAGGTGCTACATTTCGTGACACGAATGAAGATGAAATCTATGCTTTCTTTGGTATTCTGGTAATGACAGCAGTGAGAAAAGATAACCACATGTCCACAGATGACCTCTTTGATCGATCTTTGTCAATGGTGTACGTCTCTGTAATGAGTCGTGATCGTTTTGATTTTTTGATACGATGTCTTAGAATGGATGACAAAAGTATACGGCCCACACTTCGAGAAAACGATGTATTTACTCCTGTTAGAAAAATATGGGATCTCTTTATCCATCAGTGCATACAAAATTACACTCCAGGGGCTCATTTGACCATAGATGAACAGTTACTTGGTTTTAGAGGACGGTGTCCGTTTAGGATGTATATCCCAAACAAGCCAAGTAAGTATGGAATAAAAATCCTCATGATGTGTGACAGTGGTACGAAGTATATGATAAATGGAATGCCTTATTTGGGAAGAGGAACACAGACCAACGGAGTACCACTCGGTGAATACTACGTGAAGGAGTTATCAAAGCCTGTGCACGGTAGTTGTCGTAATATTACGTGTGACAATTGGTTCACCTCAATCCCTTTGGCAAAAAACTTACTACAAGAACCGTATAAGTTAACCATTGTGGGAACCGTGCGATCAAACAAACGCGAGATACCGGAAGTACTGAAAAACAGTCGCTCCAGGCCAGTGGGAACATCGATGTTTTGTTTTGACGGACCCCTTACTCTCGTCTCATATAAACCGAAGCCAGCTAAGATGGTATACTTATTATCATCTTGTGATGAGGATGCTTCTATCAACGAAAGTACCGGTAAACCGCAAATGGTTATGTATTATAATCAAACTAAAGGCGGAGTGGACACGCTAGACCAAATGTGTTCTGTGATGACCTGCAGTAGGAAGACGAATAGGTGGCCTATGGCATTATTGTACGGAATGATAAACATTGCCTGCATAAATTCTTTTATTATATACAGCCATAATGTCAGTAGCAAGGGAGAAAAGGTTCAAAGTCGCAAAAAATTTATGAGAAACCTTTACATGAGCCTGACGTCATCGTTTATGCGTAAGCGTTTAGAAGCTCCTACTTTGAAGAGATATTTGCGCGATAATATCTCTAATATTTTGCCAAATGAAGTGCCTGGTACATCAGATGACAGTACTGAAGAGCCAGTAATGAAAAAACGTACTTACTGTACTTACTGCCCCTCTAAAATAAGGCGAAAGGCAAATGCATCGTGCAAAAAATGCAAAAAAGTTATTTGTCGAGAGCATAATATTGATATGTGCCAAAGTTGTTTCTGA

**Supplementary Table 1.** Pupae showing transient *ZsGreen* expression following injection, and total adult survivors for the two rounds of microinjections. Note that not all adults were screened for transient expression at pupal stage.

|  |  | Pupae | | | Adults  (some not screened as pupae) | | |
| --- | --- | --- | --- | --- | --- | --- | --- |
|  |  | Male | Female | Total | Male | Female | Total |
| *Round 1* | Total | 26 | 30 | 56 | 30 | 30 | 60 |
|  | Transient expression | 15 | 10 | 25 |  |  |  |
|  | % transient expression | 57.7 | 33.3 | 44.6 |  |  |  |
| *Round 2* | Total | 17 | 14 | 31 | 20 | 20 | 40 |
|  | Transient expression | 11 | 11 | 22 |  |  |  |
|  | % transient expression | 64.7 | 78.6 | 71.0 |  |  |  |

**Supplementary Table 2.** Proportions of G2 pupae showing *ZsGreen* marker expression from a single screened oviposition. Pools H1, K2 and K3 failed to produce any G2 eggs and thus could not be assessed.

| Pool | ZsGreen pupae  (total) | ZsGreen pupae  (%) | Total pupae |
| --- | --- | --- | --- |
| A | 33 | 50.0 | 66 |
| H | - | - | - |
| K1 | 34 | 54.8 | 62 |
| K2 | - | - | - |
| K3 | - | - | - |

**Supplementary Text 2. *Piggybac* flanking sequences and Blastn results**

Line H

CACATATTTTCTGAATTCTCCGTAAATTTCACTTTATCCATCACCACTTTTTTCCAACGCTTCACGGAATCCAAACAAACTTGACAGTTGAGCGCGCGAAAGAGAAACAAACCAAACCAATGTTTAGAATGCAGGTGTGGCGCTGTCAGATGTCAAATGACGTGAGGTATTTAATTCCTTAATATATTAACTGCTAAAATTAATACATTAAGGCCAATGTCACGCCCCTCGCCTTAAACCATTCTTTTGTAAGAAAGGCAAAAAACAGACTACCTACAGACTTTTTGTCAAGACTATACAGACACGGCCTTTGAGGAAAATGGCATCCCTCTACACGACTTTTTCGTGGCCGTCAGACCCGATCAGTTGAGGCTATGTGAATTCAGACCATTTTTTAAACTGCTTGTAATTTCAGATAGGTAAGTCAAATTTTGAAAATTCTTACTCCACCTAAAAGGTCTTCTCATATATAACATGTGAGGGTTTCATGAAAAAACCACCCTTTTTACCGTAATTTTAATTAATATGAAACAGTTTTTTTAACATAACTGTTTTAAATACATGATAAAACCGCATGAATTTAAATAGCAACTTAGGGGACGTTAAGAAACCATTCCGCACCGCCCTCCGCACCACGCCCTTTCTGTCGTCTTCC

TTAA insertion site in red.

Culex quinquefasciatus strain JHB chromosome 3, Matches: 268

Culex quinquefasciatus strain JHB chromosome 2, Matches: 289

Culex quinquefasciatus strain JHB chromosome 1, Matches: 158

Line A

TCTGGTGCTTTACGAGCTTCCATTCGCTTGTGTGCTGATGGACTCAAAATTCTGCTACCTACCAGAAAGGATTACAACTACGTTCGGGATTTCCTGAACAACACAAAGATTGAATACTACAGCCATGACGATCCAGGTAAACGCCCCATGAAACAGGTCCTCCGTGGCCTGTACGACATGGATGTGAGTGTGCTGAAAGAAGAGCTCAAATCTCTTAAGTTGAACGTGATCGAAGTCTTCAAGATGACGAGACACAACAAGGACATCAAGTATCGTGATCAACTGTACCTGGTTCATCTCGAGAAAGGATCGACAACGCCGTCTGAGCTGAAAGCAGTTCGGGCTATTTTCAACATCATCGTGACTTGGGAACGTTATCGTCCAGTGCACCGTGACGTGAGACAATGTTCGAACTGCTTGCAGTTTGGACATGGTGGAAGGAACTGTTTCATCAAGAGTCGTTGTGCAACCTGCGGAGGTGAGCACAAAACACAAGCTTGCGAAACAATCAACGAAAACATCGAAGCGAAATGCTTCAATTGCGGTGGCGACCATTCTACCAAGAATCGAAGCTGTCCAAAACGTGCTGAGTTTGTGAAAATTCGGCAGCAAGCGACGACGAGGCACCAACCAAATCGTCGCAAAACACCACCAACTTTCACGGACGTGGATTTTCCTGCTTTGGCGTCACCTGGAGCGGGATCTGTTCGAGTGGTTCCAAATCTGCAGCCATTGCCGTTTAAGCAGCGGCAAAGAGTTGGAGAGAATACAACACCTCCAGGCTTCAGTCAGCAACCGAGGGAAAACCAACCAGCATTAACGGATGAAGGCAGTAGTGACCTGTTTTCACCACAAGAACTTCTGAACATTTTCATCGAGATGACAACAACACTGCGTGGGTGCAAAACTCGCCAGGAACAAGTAAGAACGCTTGGAGCATTCATCTTAAAATACAGTTCATAGTTTACTCGTTCTAATGATTTTGAATAATTCAAAGAATTTTTAATTTAGTTTTAAGTTAGGATCCACCGCCCTCCGCACCACGCCCTTTCTGTCGTCTTCACGCT

TTAA insertion site in red.

Culex quinquefasciatus strain JHB chromosome 3, Matches: 664

Culex quinquefasciatus strain JHB chromosome 2, Matches: 456

Culex quinquefasciatus strain JHB chromosome 1, Matches: 646

Line K

TTAATAATGCAATAATAAATGTTAAATTATGTATTCCAAATAGTTTACAATCGGTTTTTGCCTTCCTCACCTTAGATTGTAAGGTTGTATAGCCATTATCAA

Only PB 5’ flanking sequence identified.

TTAA insertion site in red.

No significant similarity found using NCBI Blastn
